# Dynamic Kidney Organoid Microphysiological Analysis Platform

**DOI:** 10.1101/2024.10.27.620552

**Authors:** SoonGweon Hong, Minsun Song, Tomoya Miyoshi, Ryuji Morizane, Joseph V. Bonventre, Luke P. Lee

## Abstract

Kidney organoids, replicating human development, pathology, and drug responses, are a promising model for advancing bioscience and pharmaceutical innovation. However, reproducibility, accuracy, and quantification challenges hinder their broader utility for advanced biological and pharmaceutical applications. Herein, we present a dynamic kidney organoid microphysiological analysis platform (MAP), designed to enhance organoid modeling and assays within physiologically relevant environments, thereby expanding their utility in advancing kidney physiology and pathology research. First, precise control of the dynamic microenvironment in MAP enhances the ability to fine-tune nephrogenic intricacies, facilitating high-throughput and reproducible human kidney organoid development. Also, MAP’s miniaturization of kidney organoids significantly advances pharmaceutical research by allowing for detailed analysis of entire nephron segments, which is crucial for assessing the nephrotoxicity and safety of drugs. Furthermore, the MAP’s application in disease modeling faithfully recapitulates pathological development and functions as a valuable testbed for therapeutic exploration in polycystic kidney diseases. We envision the kidney organoid MAP enhancing pharmaceutical research, standardizing processes, and improving analytics, thereby elevating the quality and utility of organoids in biology, pharmacology, precision medicine, and education.

## INTRODUCTION

Human kidneys are one of the most intricately developed organs in the body^1^. Its unique anatomical and physiological features, particularly when compared to those of animal models, underscore the critical need for more humanized models^2^. These models are essential for accurately studying human kidney development, the mechanisms of kidney diseases, and evaluating the toxicity of pharmaceuticals and environmental agents on renal function.

In this pursuit, kidney organoid models have risen to prominence as indispensable tools for biological studies. They illuminate the complex physiological mechanisms that govern lineage specification, interactions, and responses throughout the course of organ development and pathogenesis. These models hold immense promise across various domains, including renal development biology, disease modeling, biomarker discovery, drug screening, personalized medicine, and education and training. The emergence of an ideal organoid model is eagerly awaited to unlock their full potential.

Nonetheless, a well-recognized challenge in using organoids as human physiological models is the variability among organoids, even when developed under well-defined protocols. This variability can affect biological reproducibility across different timeframes and locations, creating difficulties for systemic quantitative analysis of kidney organoid changes. For an organoid model to serve as a reliable alternative to living model systems, achieving modeling consistency is essential. This consistency allows the organoid to function as a biological predictive model that demonstrates robust responses to external stimuli, enabling systematic investigation. While intrinsic cellular properties, such as iPSC differentiation potential, can pose challenges in establishing reproducible predictive organoid models, it is equally crucial to recognize that much of the variability arises from extracellular factors linked to conventional static culture conditions. In addition, the skill and expertise of researchers involved in organoid cultures can exert a substantial influence on outcomes due to inconsistent manual manipulation and handling. It is noteworthy that this particular concern can be effectively addressed through a bioengineering approach, as demonstrated in this study.

The development of the kidney, orchestrated through reciprocal signaling, entails a complex web of interactions between different cell types and molecular signals^3^. A diverse array of molecules partakes in this intricate process, with specific factors being key drivers and other components allowing for precise adjustments in development. Notably, there is currently no method available to generate single renal populations and reconstruct a kidney’s architecture by assembling individual renal populations. This underscores the vital role of heterotypic cell-to-cell dynamic interaction, which remains incompletely understood in current renal biology, in generating renal populations in vitro. Especially, extracellular signaling during human kidney development occurs through molecular secretion and receptor signaling among nearby populations^4-7^, therefore, the technique of endowing well-regulated environments becomes critical in refining organoid modeling. While the “minimal factor” approach to organoid differentiation is commonly used, it often overlooks the balance between spontaneous intercellular communication and exogenous differentiation factors in current protocols^8-11^. This oversight may contribute to the unpredictability of concurrent organoid modeling. Addressing the balance between external minimal factors and intercellular communication within the extracellular space is essential for refining organoid models to more accurately replicate kidney development with better reproducibility.

Herein, we introduce a kidney organoid microphysiological platform (MAP) designed explicitly for generating uniform kidney organoids and facilitating organoid assays (Fig. 1). To address organoid heterogeneity and poor reproducibility caused by inadequate regulation of biochemical extracellular space, our kidney organoid MAP is designed to create biochemically regulated extracellular environments through microfluidic perfusion dynamics. The MAP’s interstitial spaces are structured as small-volume, microliter-scale hemispherical chambers to enable a reproducible and reliable miniaturization of organoids through ensuring a balance of intrinsic and extrinsic factors. This design minimizes stochastic variability observed in larger tissue constructs, thereby promoting more controlled and consistent organogenesis compared to traditional culture methods. The hemispherical chambers arranged along identical perfusion channels promote high biochemical similarity across the organoid array, supporting robust statistical analysis to enhance the understanding of biological significance.

**Figure 1.**
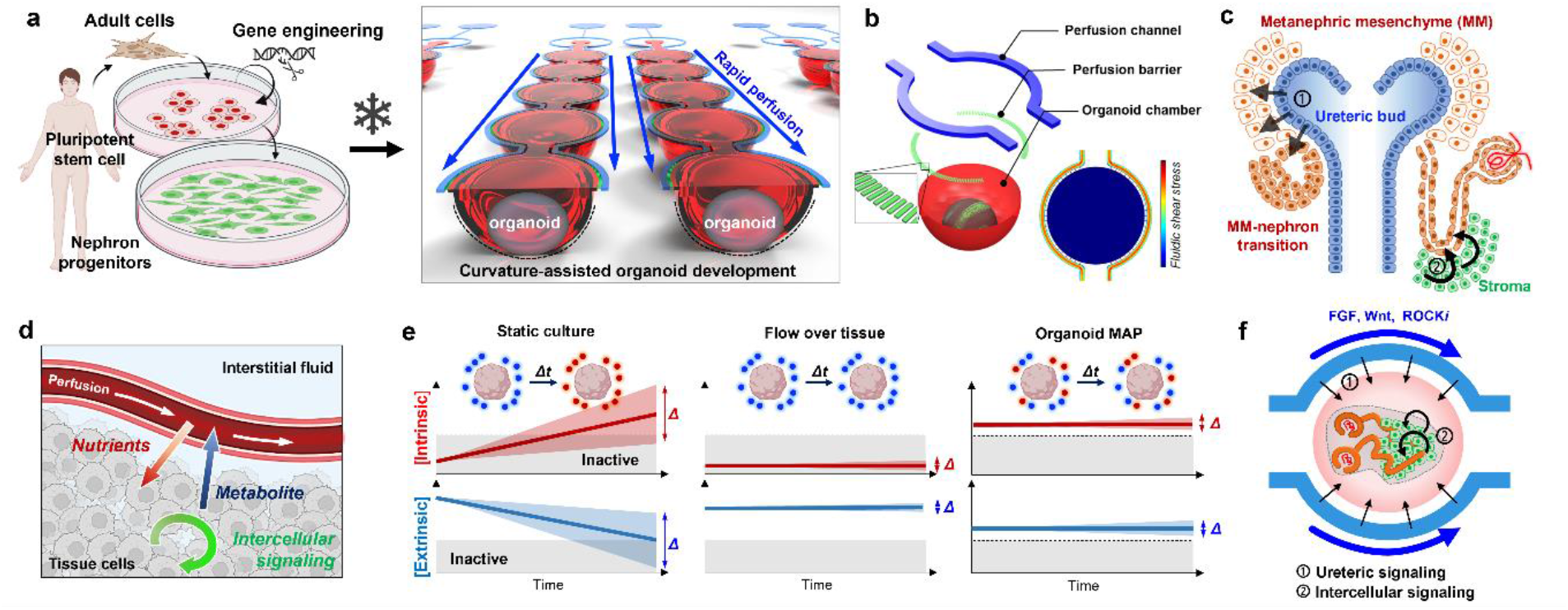
Kidney Organoid MAP for Reproducible Organoid Modeling. **a**, The on-chip kidney organoid generation process involves a sequential procedure, starting with the generation of nephron progenitor cells (NPCs) in large-scale 2D culture, followed by the intermediate cryopreservation of NPC aliquots, and culminating in dynamic perfusion organoid development within the MAP. **b**, The functional microfluidic unit of the organoid MAP is comprised of perfusion channels, a hemispherical organoid chamber, and interconnecting perfusion barrier nanochannels. This fluidic configuration enables biomimetic biomolecular interactions across three compartments without subjecting the organoid chamber to fluidic shear stress. **c**, In vivo nephrogenesis of metanephric mesenchyme into polarized nephrons occurring through signals from the ureteric epithelium and stromal cells. **d**, In vivo fluidic circulation mimicked in the MAP’s microfluidic functional unit, allowing the separation of fluid momentum and functional tissue cells through the barrier properties of the endothelium lining. **e**, Comparison of three organoid culture methods highlighting the delicate balance within the MAP of intrinsic and extrinsic factors in close proximity to cells over time. In contrast, static culture conditions experience inevitable changes in these factors over time due to intermittent changes in media. Flow over tissue culture, on the other hand, is unsuitable for accumulating spontaneous cellular secretions around cells, which play a critical role in current organoid protocols. The symbol Δ represents uncontrollable variations in biomolecules, contributing to developmental variability in organoid modeling. **f**, MAP’s signaling to facilitate homogenous, reproducible development of kidney organoids by implementing dynamic microenvironments for the ureteric bud signaling (FGF, Wnt), fostering progenitor proliferation including stromal cells (ROCK inhibition), and regulating controlled intercellular communication (stroma-tubule interaction) within the MAP’s interstitial environment.

Our MAP’s generation of kidney organoids demonstrates its potential to enhanced reproducibility, controllability, and robust miniaturization of the physiological human model derived from human stem cells. The well-regulated biochemical environment in MAP enables precise control over the developmental fates of kidney populations, allowing for the accurate specification of nephron details between proximal and distal segments. The organoid reproducibility in MAP facilitates systemic iterative exploration of molecular signaling during early kidney development, and the MAP’s organoid miniaturization advances a precise quantitative assessment of entire nephron structures. Utilization of the MAP’s uniform and miniaturized organoid modeling and perfusion dynamics has proven valuable for nephrotoxicity studies, establishing it as a powerful tool for pharmaceutical applications. Additionally, the application of genetic engineering in disease modeling within MAP enables a systemic approach to therapeutic discoveries. The potential of MAP, as demonstrated here with kidney organoid modeling, leads us to believe that it will play a crucial role in advancing organoid models closer to an ideal in vitro yet in vivo-like human physiological model.

## RESULTS

### Kidney organoid MAP designed for on-chip organogenesis

To provide precise biochemical control and maintain consistent metabolic conditions (including extracellular nutrient and metabolite levels), our kidney organoid MAP is engineered within a microfluidic framework that enables continuous perfusion around the organoid arrays (Fig. 1a). The MAP’s fluidic dynamics was designed to prevent fluidic shear stress on cells similar to the in vivo perfusion and to create a balanced interplay between extrinsic differentiation factors and intrinsic cellular secretion within the vicinity of the cells (Fig. 1d). Additionally, the space where the organoid is positioned was designed as a capacious concave-bottomed chamber to enable unimpeded tissue growth while ensuring a central location for optical tracing purposes. These considerations resulted in a functional unit comprising an array of hemispherical chambers, bifurcated perfusion channels, and interconnecting nanochannels designed to emulate the properties of an endothelium barrier (Fig. 1b). The overall layout comprises eight functional units, with each unit housing a ten-organoid array, surrounded by perfusion channels connected to the same media reservoirs. As a result, a single kidney organoid MAP accommodates 80 organoids, which are divided into eight groups, each exposed to distinct biochemical perfusion conditions, facilitating statistical analysis based on ten organoids per group.

The MAP fluidic design creates a buffering system with a time constant of several hours, enabling a well-balanced, temporally stable equilibrium of intrinsic secretions and extrinsic factors around the organoid (Fig. 1e). This stands in stark contrast to conventional configurations, such as static culture with intermittent media changes^8-11^ and direct flow over organoids^12,13^, which resulted in an inadequate balance between intrinsic and extrinsic substances, lacking temporal dynamics and control. Given the significance of fluidic dynamics in our MAP’s biological modeling and streamlined experimental process, we have implemented a tubeless fluidic perfusion system that operates automatically through hydraulic pressure-driven flow between media reservoirs. This system is enhanced by a custom-made tilting plate controlled by a microcontroller and a motorized linear actuator, incorporating feedback from an angular sensor to maintain a consistent flow rate over time. This setup facilitates accurate microfluidic operations over a month-long organogenesis program, substantially reducing the complexity and minimizing the operational challenges typically associated with fluidics.

### MAP enables kidney organoid development under biochemically defined perfusion dynamics

To enhance the reproducibility of kidney organoid modeling, the on-chip organogenesis in MAP begins with the reaggregation of SIX2-positive nephron progenitor cells (NPC) derived from a large-scale pool cryopreserved in separate aliquots. For this, first, we differentiated human pluripotent stem cells into metanephric mesenchyme in a monolayer by following our previous protocol^14^, which included the timely application of CHIR99021 (to active Wnt/β-catenin signaling), activin, and FGF9. At the end of this stage, we prepared a large-scale pool of NPCs through single-cell dissociation and resuspension, subsequently cryopreserving them in 2 million cell aliquots. The availability of ample cryopreserved NPC stocks facilitated the fine-tuning of MAP culture protocols and enhanced the reproducibility of kidney organoids, even in experiments conducted at different times.

Then, we meticulously programmed the perfusion dynamics and biochemical composition within MAP fluidics to guide nephron progenitors toward the development of nephron-enriched organoids. In our programming, MAP’s perfusion serves as a steady, temporally stable source, effectively emulating ureteric bud signaling to metanephric mesenchyme, fostering the formation of pretubular aggregates. Simultaneously, the fluidic, motion-free, yet molecule-permeable interstitial chamber provides microenvironments conducive to clear-cut intercellular communication, a crucial factor for spatially patterning nephron segments (Fig. 1c & 1f). Furthermore, MAP’s timely regulation of Rho signaling provides highly effective control over the abundance of distal segments and stromal cells within organoids, underscoring MAP’s modeling precision.

First, the initial formation of NPC spheroids within the MAP was optimized to enhance cell viability and support organoid development into miniaturized functional tissues. A brief 8-hour treatment with a ROCK inhibitor (Y-27632) at the beginning of MAP’s NPC culture significantly improved cell viability and spheroid formation, with a single spheroid forming in each chamber within 2 to 4 hours, facilitated by the hemispherical curvature. During the subsequent 2-day CHIR99021 priming, the organoid development in MAP underscored the importance of secretory signaling in guiding NPCs toward metanephric condensation. During this stage, the flow rate was set to minimize the washout of autocrine/paracrine factors around the cells; otherwise, progress in organogenesis could be hindered (Fig. 2a). Subsequent three-day developmental conditions with FGF9 supplementation, maintained at the baseline perfusion rate, guided all spheroidal tissues along a consistent morphological development trajectory. After reaching the renal vesicle stage, an additional week with basal medium flow led to minimal morphological and volumetric changes over the subsequent four weeks, providing ample time for assessing organoid responses to pharmaceutical compounds.

**Figure 2:**
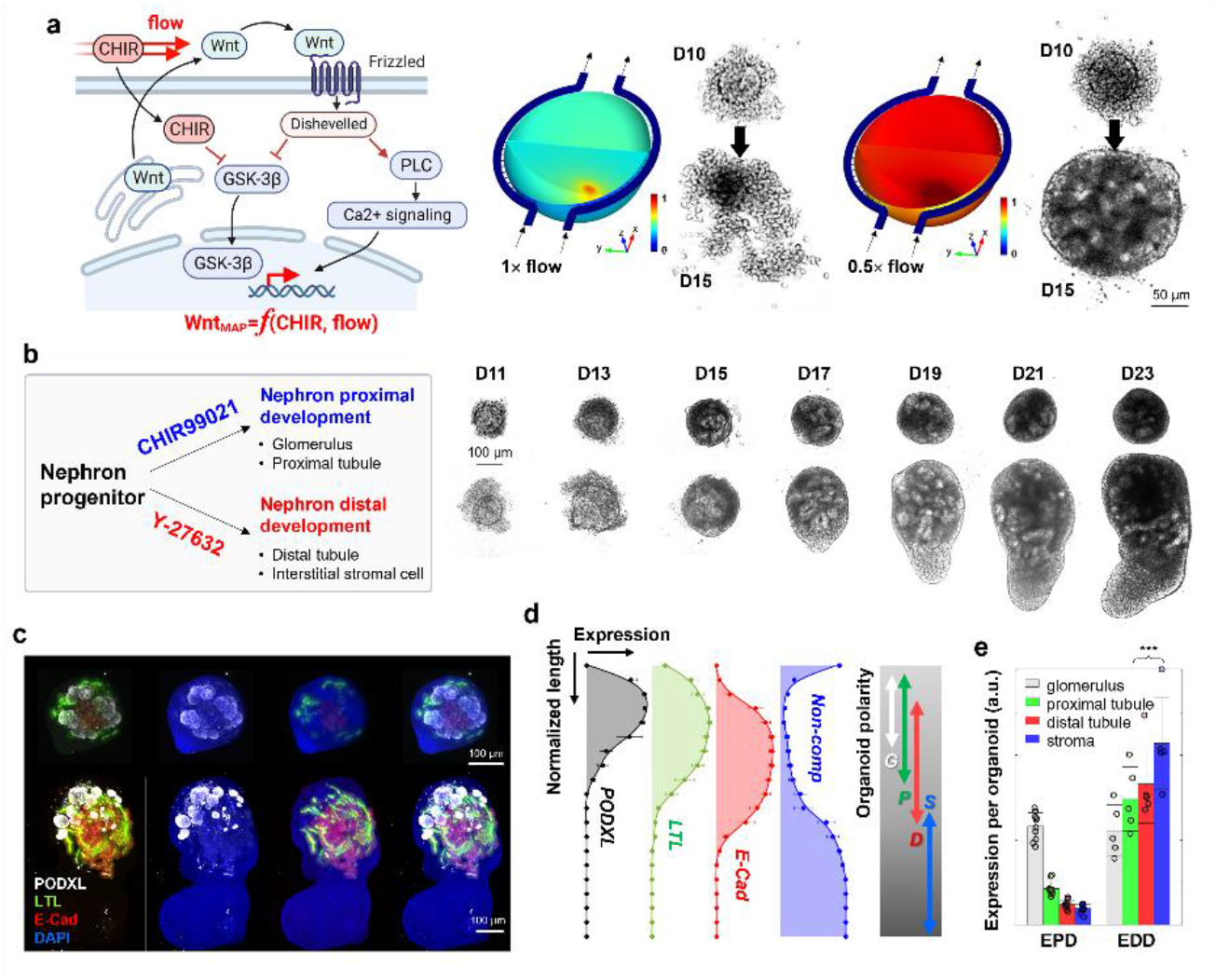
Biochemical Precision and Controllability of MAP’s Organogenesis. **a**, The initial step of the MAP organogenesis, involving CHIR99021 priming of NPCs, emphasizes the crucial role of secondary signaling in addition to canonical Wnt/β-catenin signaling. The dynamic flow within MAP demonstrates that this signaling occurs in the context of secretion and receptor binding, functioning in an autocrine/paracrine manner. Computational and experimental results underscore the importance of secretion molecules, independent of Wnt/β-catenin easily governed by CHIR99201 perfusion, in facilitating the transition of NPCs into early epithelial nephron constructs. **b**, MAP’s precision in NPC differentiation signaling allows for the biochemical modulation of organoid development into either proximal-enriched or distal-enriched nephron development through the timely application of CHIR99021 and Y-27632. Optical monitoring within MAP exhibits the distinct developmental trajectories of these two types. Both patterns share a similar initial phase in NPC nephrogenesis, while tubules and stromal cells thrive in the later phases of the enhanced distal development. **c**, Structural analysis using confocal microscopy reveals the characteristic features of the proximal and the distal enrichment, especially significant distinction of distal tubules and stromal cells. The labels PODXL, LTL, ECAD, and DAPI correspond to podocalyxin, lotus tetragonolobus lectin, E-cadherin, and 4’,6-diamidino-2-phenylindole, respectively. **d**, The structural arrangement of MAP’s organoids with enhanced distal development reflects a sequential placement from PODXL-positive glomerulus-like structures, proximal tubules, and distal tubules with stroma, resembling early-stage kidney arrangement (Data are represented as the mean ± s.d. from 5 organoids in three independent experiments; data points excluded for visual clarity). **e**, Quantification of individual compartments indicates the prevalence of E-CAD positive distal tubules and the stromal population in enhanced distal development (EDD) compared to enhanced proximal development (EPD) (n_EPD_ =10 and n_EDD_ =5 from two and three independent experiments, respectively, and presented as the mean ± s.d). The normalized expression levels of the proximal tubule, distal tubule, and stroma showed a significance level in the *t*-test of *p* < 0.01 (***).

Using the NPC cryopreservation pool, we iterated the aforementioned organogenesis steps while controlling the initial cell numbers to ensure the consistent generation of miniaturized kidney constructs. MAP’s cell loading concentration, ranging from 0.2 to 10 million cells per milliliter (MPM), led to the reproducible development of kidney organoids with proportional overall and glomerular volumes. Initial cell concentrations below 0.2 MPM were associated with a higher rate of organoid development failure, such as lack of proliferation or epithelialization, indicating a minimum cell threshold necessary for successful nephrogenesis in MAP’s biochemical conditioning. Following the established organoid protocol^11^, the on-chip organogenesis in the MAP consistently resulted in spherical tissues predominantly composed of proximal segments with limited distal tubular expression, irrespective of the initial cell number and organoid size. The organoids maintained morphological stability for 1.5 months under the MAP’s dynamic culture conditions, without significant cell deaths and morphological changes. Additionally, the MAP organoids exhibited de novo endothelial vascularization, emerging near the glomerulus-like structures.

### MAP organogenesis accurately fine-tunes kidney development in organoids

Application of the established protocol within MAP consistently yielded organoid patterns enriched with nephron-proximal segments, such as the Podocalyxin (POXDL)-enriched glomerulus-like structures and LTL-positive proximal tubules. This result indicates that the continuous balanced biochemical refreshment provided by MAP during CHIR99021 priming more prominently directed the developmental fate of NPCs towards nephron proximal segments, compared to organoids in static culture^11^.

Moreover, our kidney organoid in MAP exhibited precise control over proximal-distal development with modulation of ROCK inhibition (Fig. 2b), aligned with prior studies showing that timely ROCK inhibition supports NPC and stromal proliferation^15,16^. When we extended the treatment duration of Y-27632 by an additional 2 days, coinciding with CHIR99021 priming, the subsequent organoid development underwent a dramatic alteration, characterized by enhanced expression of distal tubular segments and stromal compartment. The time-lapse monitoring within MAP demonstrated that this distinctive development consistently occurred after completing extrinsic FGF9 perfusion and persisted for ten days with gradual morphological changes. This developmental modulation by ROCK inhibition was not obviously observed under static culture conditions, highlighting the more refined modulation of organogenesis achievable in the dynamic MAP environment.

In confocal imaging, the organoids with enhanced distal development (EDD) exhibited larger proportions of E-Cadherin-positive distal tubules compared to the organoids with enhanced proximal development (EPD) (Figs. 2c & 2e). Furthermore, the EDD organoids displayed a consistent structural arrangement along a pseudo-axis, involving sequential positioning of the glomeruli, proximal tubules, distal tubules, and stroma-dominated regions. This arrangement resembled that of an early-stage kidney^17^, with the cortex enriched in epithelialized compartments and a non-epithelialized medullary region. As such, we defined this arrangement along a pseudo-corticomedullary axis, and our quantification of nephron biomarkers supported the inclination for nephron arrangement during organoid maturation in MAP (Fig. 2d), highlighting intriguing interactions among kidney lineages^18^.

### Kidney organoid MAP facilitates quantitative nephrotoxicity study

The reproducibility and homogeneity of kidney organoid models allow for comprehensive, detailed characterization of drugs’ nephrotoxic properties within MAP. The ability to precisely address individual organoids, the intentional miniaturization of kidney organoids, and on-chip whole-mount confocal microscopy enable thorough examination of nephron segment-specific damage within each organoid. To further enhance analytics, we developed an evaluation method that assesses nephrotoxicity by dividing whole-mount confocal images into micro-quantized segments and calculating the percentage of segments displaying specific signs of nephrotoxicity (Fig. 3a). In a proof-of-concept demonstration, we assessed the effects of conventional drugs, gentamicin^19^ and cisplatin^20^. These drugs are commonly recognized as nephrotoxic but are still in use due to their effectiveness and the lack of alternatives. Therefore, a precise understanding of their nephrotoxicity may aid in mitigating side effects.

**Figure 3.**
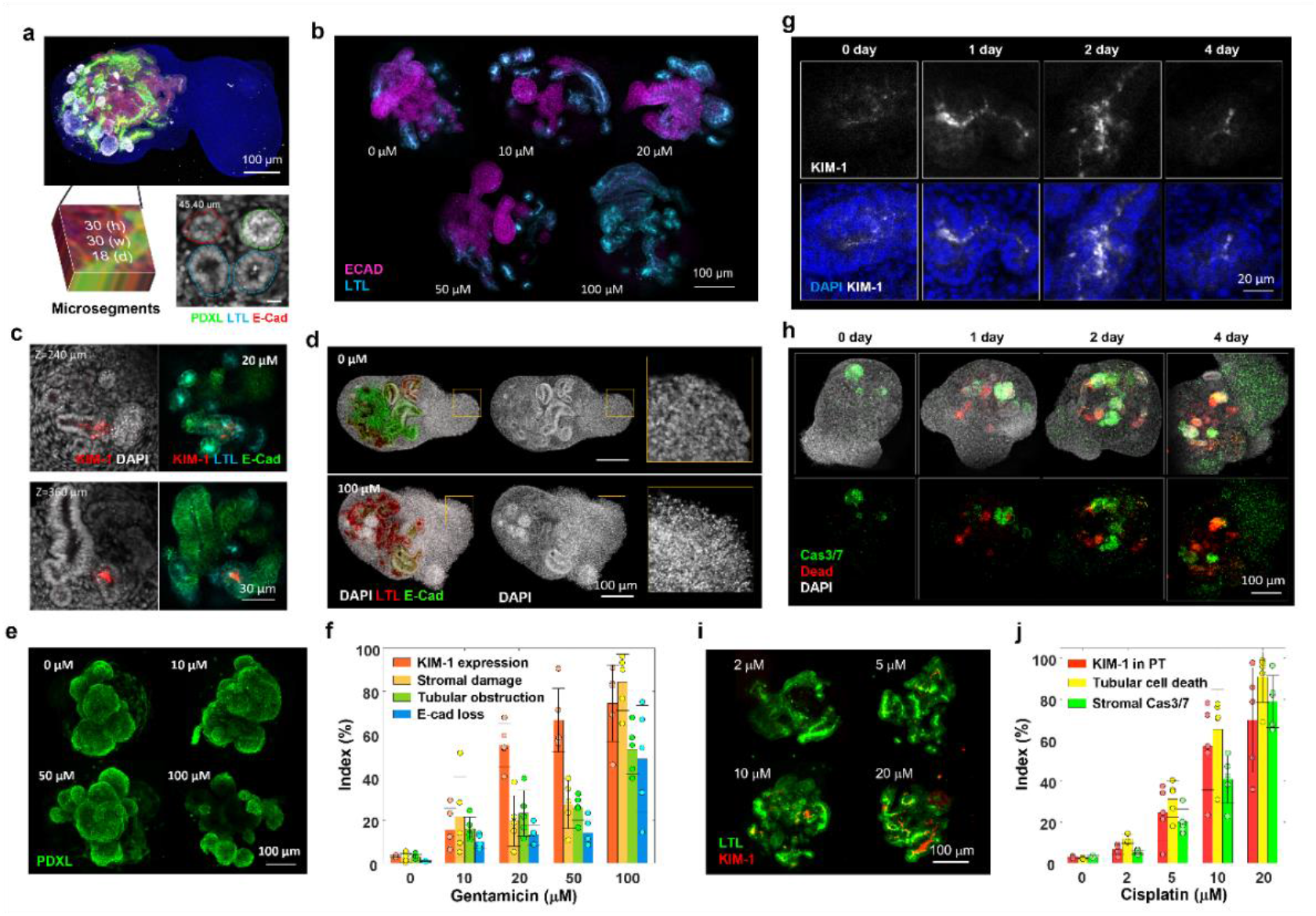
MAP’s nephrotoxicity study with nephron segment-specific quantification. **a**, MAP’s nephrotoxicity quantification is facilitated by MAP’s fidelity in organoid miniaturization, whole-organoid confocal microscopy, and microsegment quantification. Our optimal compromise between microscopic visualization of structural details and specimen size allows effective 3-dimensional reconstruction and a multi-context analysis via microsegmental examination. A snapshot presents the optical power to detail structural contexts. **b, c, d, e**, 2-day-long gentamicin-containing perfusions in MAP causes a broad aminoglycoside nephrotoxicity across nephron constructs (day 30). This study included an examination of distal tubule damage with decreased E-CAD expression (b), a tubular injury marker of proximal injury that increases KIM-1 (c), tubular obstruction with nuclear staining (the inset in a), cell death among stromal population by nuclear morphology (d), and glomerular damage by reduced podocalyxin expression (e). LTL-based proximal tubular damage was not clearly distinguished. **f**, The dose-dependent gentamicin nephrotoxicity is summarized in a bar plot on the multi-aspect nephrotoxicity through the microsegmentation approach. Among the broad spectrum, KIM-1 expression was the most sensitive biomarker induced by gentamicin. Most comparisons to the control showed statistical significance in the *t*-test (*p* < 0.05), except for KIM-1 at 10 µM gentamicin, which had a *p*-value of 0.052, as indicated in the raw data. **g, h**, Temporal dynamics of kidney damage expression after 12-hour 10-µM cisplatin perfusion were characterized in time-lapse application of assays across a uniform set of kidney organoids (day 30) in MAP. KIM-1 in the lumen of proximal tubule, and appearance of apoptotic and dead cells gradually increased after the cisplatin application. Our analysis presents a constituent expression of caspase 3/7 activity in the PODXL-positive compartment, requiring caution to interpret the expression per organoid. **i, j**, Cisplatin dose-response was quantified 2 days after 12-hour cisplatin perfusion with concentration between 2 and 20µM. The quantification of KIM-1 and dead cells was specified within LTL-positive proximal tubules. The caspase-active apoptotic cells rarely overlapped with tubular segments but increased its prevalence among stromal cells. The zero concentration represents a vehicle control with DMSO used to prepare the cisplatin stock. All the comparisons to the control showed statistical significance in the *t*-test (*p* < 0.05), as indicated in the raw data. The nephrotoxicity plots of **f** & **j** were derived from distinct experiments, comprising seven for gentamicin and three for cisplatin. Each data is the mean ± s.d of 5 organoids per condition.

In the gentamicin nephrotoxicity study, we evaluated its broad nephrotoxic effects and a range of gentamicin dosages on the integrity of the glomerulus, proximal tubules, distal tubules, and stroma using immunostaining with anti-podocalyxin antibody (Ab), lotus tetragonolobus lectin (LTL), anti-E-Cadherin Ab, and anti-Kidney injury molecule-1 (KIM-1) Ab (Fig. 3b to 3e). Additionally, we analyzed nuclear distribution, which was valuable for assessing stromal cells and tubular obstruction. In our toxicity characterization, the extent of LTL staining remained largely unchanged after drug applications, making LTL staining a useful tool for distinguishing proximal tubules among tubular arrangements. To the contrary, KIM-1 expression highlighted the sensitivity to identify proximal tubular injury, even when the brush border of the proximal tubule is preserved.

Our analysis of micro-segments within a 3D organoid reconstruction the effective assessment of nephrotoxicity metrics associated with gentamicin. As shown in the comparative graph in Fig. 3f, the analysis indicated that KIM-1 expression in LTL-positive proximal tubules was particularly sensitive to increases in gentamicin dosage. Stromal damage and tubular obstruction also increased in a dosage-dependent manner. E-cadherin loss, indicative of epithelial integrity, remained minimal except at the highest gentamicin concentration (100 μM), where significant loss indicated substantial distal tubule injury. Podocyte damage as reflected by podocalyxin loss was not clearly observed in our dose-response study, consistent with findings from other studies^19^.

In the cisplatin study, MAP’s organoids provided an effective model for characterizing the toxicokinetic processes of acute cisplatin nephrotoxicity, offering consistent and reproducible organoid arrays. Focusing on the proximal tubule, a well-known target of cisplatin toxicity, we characterized the time course of KIM-1 expression following a 12-hour perfusion with 10 μM cisplatin, during a subsequent cisplatin-free recovery period lasting from 0 to 4 days (Fig. 3g). Confocal imaging of organoids fixed at specific time points revealed the dynamics of KIM-1 accumulation and dissipation within the tubular lumen, with peak expression observed on day 2 post-cisplatin treatment, rather than at earlier time points. We also examined experimental replicates to assess apoptotic cells using a caspase 3/7 assay and cell death with membrane-impermeable propidium iodide (PI) (Fig.3h). The image analysis showed that the number of PI-positive cells were mainly found in the tubular segments and gradually increased over time starting from day 1 after cisplatin exposure. However, the caspase 3/7-positive regions were distinct from the PI-positive regions, particularly as they were not observed in the proximal tubules. This aligns with that necrosis is a key mechanism underlying the loss of proximal tubular cells by the physiological cisplatin dosage^21^. In contrast, caspase activity was scattered within the stromal population and concentrated in the glomeruli. The gradual increase in caspase 3/7-positive stromal cells suggests a distinct mechanism of stromal nephrotoxicity compared to that observed in the proximal tubule. In the case of glomerular caspase 3/7 positivity, elevated caspase activity was also observed in the glomeruli of untreated control organoids, which may reflect natural processes occurring during glomerular development in kidney organoids.

In the subsequent dose-response experiments, we set the endpoint to day 2 post-cisplatin treatment to characterize nephrotoxicity through KIM-1 expression in the proximal tubule lumens, caspase 3/7 activity in the stroma, and cell death in the proximal tubules (Fig. 3i & 3j). Quantification of these metrics showed a clear dosage-dependent response. KIM-1 expression and proximal tubular cell death exhibited similar dosage-dependent profiles, manifesting the effectiveness of KIM-1 as a biomarker for proximal tubular damage. Yet, analysis of the perfusates throughout the post-cisplatin recovery period revealed no detectable levels of soluble KIM-1 in the media (LOD ∼ 0.5 ng/mL) across all doses, suggesting that soluble KIM-1 may not be a suitable biomarker in organoid toxicity assays. Organoids exposed to concentrations higher than 10 μM cisplatin showed over 50% of proximal tubular segments with both cell death and KIM-1 expression. At those dosages, caspase 3/7 activity was also significantly detected in the stromal population, indicating that cisplatin-induced nephrotoxicity affects both proximal tubule and stromal cells through distinct cellular damage responses at physiologically relevant concentrations^22^. Overall, this proof-of-concept nephrotoxicity study demonstrates that systemic toxicological quantification can be effectively achieved using organoids comparable across large-scale batches of MAP’s physiological modeling.

### PKHD1-mutant kidney organoid in MAP: replicating ARPKD pathogenesis

Among genes associated with polycystic kidney disease, a mutation in the PKHD1 gene leads to early pathological progression of the disease, causing an autosomal recessive form known as autosomal recessive polycystic kidney disease (ARPKD), which is often apparent in infancy or early childhood ^23^. To study this disease using organoid modeling, we created a PKHD1 mutant stem cell line through CRISPR/Cas9 gene editing (as previously described^24^) and compared organoid development with isogenic control organoids in MAP. While insignificant differences were observed in earlier stages before NPCs appeared, it was noticeable that the PKHD1-mutant NPCs exhibited poor viability and aggregation at the beginning of MAP culture. This necessitated an increase in the initial cell number, for size-matched initial aggregation, by a factor of approximately 1.5 for the PKHD1-mutant organoids (hereafter referred to as ARPKD organoids). Additionally, the two-week MAP organogenesis resulted in organoids that remained relatively smaller than the control even after the initial size matching.

During the MAP organogenesis periods, approximately half of the ARPKD organoids (55±4.5%) exhibited spontaneous formation of cyst-like compartments along their natural developmental trajectory (Fig. 4c). These cystic structures primarily involved E-cadherin-positive epithelial cells with a small proportion of interspersed LTL-positive cells. Most of the cyst-like compartments were attached to tubules within the organoids. Also, secondary buds from the cystic compartments and completely detached cystic structures were often observed among the organoids. In quantifying cystogenesis within this model, we found that the ratio of organoids with cystic compartments was a more reliable metric than measuring cystic volumes, as cysts exhibited fluctuating volumes but rarely disappeared once formed. Therefore, spontaneous cystogenic morphogenesis, combined with measuring the ratio of cystic organoids in ARPKD arrays within the MAP, provides an effective method for studying pathogenesis and conducting pharmaceutical screening.

**Figure 4.**
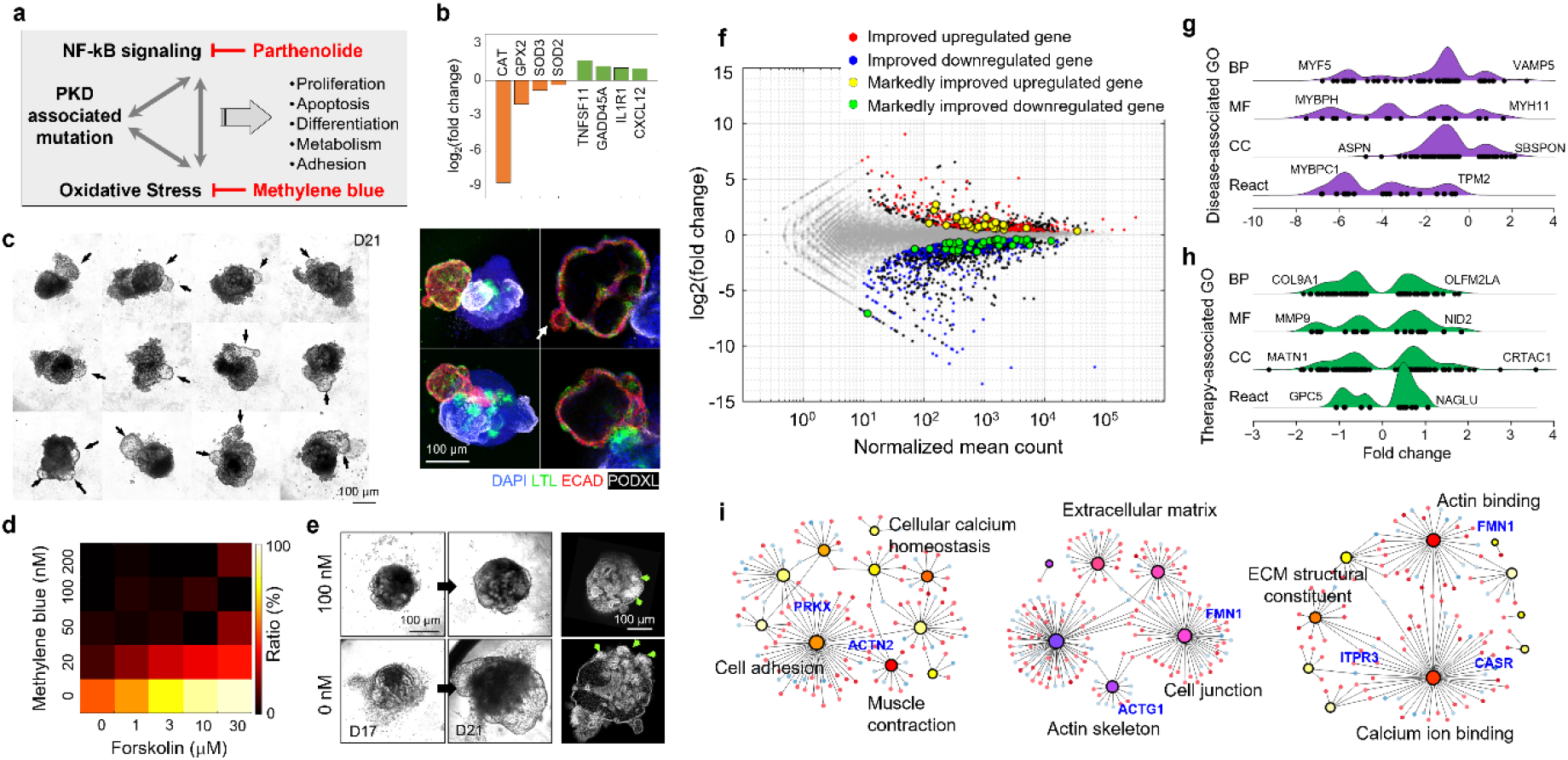
MAP’s disease modeling and therapeutic investigation based on nephrogenesis and transcriptomic expression of isogenic healthy and PKHD1-mutant organoids to recapitulate autosomal recessive polycystic kidney disease (ARPKD). **a**, The framework of our MAP’s therapeutic interventions in ARPKD pathogenesis focuses on NF-κB and oxidative stress and the use of minimal-adverse-effect drugs. Parthenolide and methylene blue were employed to regulate NF-κB and reactive oxidative stress during the early-stage cystogenesis of ARPKD. **b**, Differential expressions in ARPKD organoids include genes related to endogenous ROS scavengers and NF-κB regulation, illustrating the pathogenetic paradigm. **c**, The ARPKD organoids exhibited spontaneous development of cystic compartments with a 55±4.5% prevalence, as indicated by the arrow. These compartments were predominantly lined with E-Cadherin-positive epithelial cells, with interspersed LTL-positive cells. Additionally, secondary cyst buds were frequently observed, as denoted by the white arrow in the right confocal image. **d**, The therapeutic effectiveness of anti-ROS and NF-κB inhibitory agents was evaluated under conditions inducing cAMP-enhanced cystogenesis with forskolin. The heatmap on the left illustrates the frequency of cyst development within 20 ARPKD organoids. Parthenolide was administered with a one-day-long application. Methylene blue was administered for eight days at various concentrations prior to two-day-long Forskolin applications. **e**, The time-lapse trace presents a significant expansion of cystic compartments by forskolin application, while the therapeutic pretreatment effectively prevented pathological progression (cases depicted for 0 and 100 nM methylene blue with 500nM parthenolide prior to 30 µM forskolin). The corresponding confocal characterization with DAPI (right upper and lower panels) highlighted the exacerbation of pathological changes by cAMP signaling, whereas the therapeutic pre-treatment resulted in a normal development, comparable to their healthy organoid counterparts. The green arrows, which indicate glomerular positions, demonstrate the abnormal placement of glomeruli in cyst-containing ARPKD organoids. **f**, RNA-Seq comparisons among isogenic healthy, untreated ARPKD, and treated ARPK organoids are visualized in the baseline MA plot of healthy organoid vs. untreated ARPKD. The gray dots represent genes identified through the analysis of differential gene expression, while the black dots denote genes with a significance level of adjusted p<0.05. In comparison between untreated and treated ARPKD organoids, therapeutically improved genes among black-dotted genes were marked with the colored symbols as outlined in the legend. Additional details can be found in ‘Results,’ and ‘Methods.’ **g**, The gene ontology of the differentially expressed genes (DEG) with significance reveals PKHD1-induced modified genes that are related to skeletal muscle tissue development in biological process (BP), structural constituent of muscle in molecular function (MF), proteinaceous extracellular matrix in cellular component (CC), and striated muscle contraction in reactome (Re). Gene symbols mark the most significantly differentially expressed genes in each context. The full gene set is shown in Supplementary Table 1. **h**, The gene ontology of the DEG between untreated and treated ARPKD organoids highlights extracellular structure organization in BP, collagen binding in MF, proteinaceous extracellular matrix in CC, and HS-GAG degradation in Re. The complete gene set is provided in Supplementary Table 2. **i**, The gene set associated with therapeutic improvement was analyzed in the network analysis, resulting in an exploration of PKD-associated biology with the highlighting of blue-colored genes. The criterion applied was the significance of differentially expressed genes (DEGs) between untreated and treated ARPKD, along with the opposite fold changes between control vs. untreated ARPKD and untreated vs. treated ARPKD. Full gene names are provided in Supplementary Fig. 1.

### MAP’s ARPKD study demonstrates the efficacy of ROS and NF-κB modulation in addressing cystogenic pathogenesis

Having successfully recapitulated the pathological phenotype, we sought to characterize potential strategies for mitigating cystogenesis by controlling reactive oxygen species (ROS)^25-28^ and NF-κB signaling^29-32^ (Fig. 4a). In a proof-of-concept study for this therapeutic hypothesis, we tested the efficacy of parthenolide as an NF-κB inhibitor and methylene blue (MB) as a ROS scavenger in preventing cystogenesis in the ARPKD organoids. For parthenolide, we aimed for a short-duration, low-concentration application, considering that NF-κB is an essential signaling pathway in nephrogenesis^33^. The application of parthenolide was the most effective before pretubular aggregation, compared to other scheduled additions or no-treatment control. A low concentration of parthenolide, specifically 500 nM, was sufficient for its effectiveness, but longer applications than the one-day duration had adverse effects on organoid growth and viability. In contrast, the MB treatment could be extended due to its lower cytotoxicity, with various concentrations tested between day 9 and 16, resulting in a concentration-dependent reduction of cystogenesis. As shown in Fig. 4d, a seven-day treatment with 50 nM MB led to a developmental improvement, reducing the proportion of the ARPKD organoids with cystic compartments from 50% to 5%.

Our therapeutic approach also demonstrated effectiveness in preventing facilitated cystogenesis driven by increased cAMP signaling (Fig. 4d). Given that intracellular cAMP is a promoting factor in PKD pathogenesis, the cystogenesis of ARPKD organoids exhibited a concentration-dependent response to forskolin—a potent agonist of adenylyl cyclase known to elevate intracellular cAMP level, surpassing the baseline cystogenesis observed in the untreated culture. Two-day-long treatment of 10 µM or higher forskolin rapidly induced the development of cystic compartments in most ARPKD organoids, with volumes larger than those in untreated ARPKD organoids (>90%). MB pretreatment before forskolin significantly restrained cystic development. The propensity for cystogenesis, which was over 90% with 30 µM forskolin alone, was reduced to less than 15% with the therapeutic pretreatment (Fig. 4e).

To elucidate the transcriptomic response involved in pathogenesis and the therapeutic effect, we compared the total intracellular RNA profiles of isogenic normal organoids, untreated ARPKD organoids, and ARPKD organoids with therapeutic treatment (Fig. 4f). Alongside substantial morphological variations, ARPKD organoids exhibited significant alterations in 2,496 genes, with 875 upregulations and 1,621 downregulations, compared to the normal organoids. Notably, among the genes within the focused pathogenetic features, we observed significant downregulation of CAT (which encodes catalase, a crucial endogenous ROS scavenger) and upregulation of TNFSF11 (which encodes a member of the tumor necrosis factor) and GADD45A (a gene involved in apoptosis and microtubule stability) (Fig. 4b). In gene ontology (GO) analysis, the altered gene set was categorized into contexts related to muscle structure and extracellular matrix, presenting a coherent biological narrative to PKD studies^34-36^ (Fig. 4g).

The therapeutic treatment induced a similar number of genes (2,453) in differentiated gene expression between the untreated and the treated ARPKD organoids. This gene set exhibited GO contexts primarily associated with extracellular matrix organization, with a specific emphasis on collagen binding (Fig. 4h). The 771 genes associated with therapeutic improvement revealed a more precise impact on biological contexts, including cell adhesion, junction formation, extracellular matrix organization, actin skeleton regulation, and calcium ion binding (Fig. 4i). All the contexts have been addressed in PKD-related pathogenesis, highlighting the effectiveness of the proposed therapeutic framework validated through organoid modeling in MAP.

## DISCUSSION

In this article, we introduce a meticulously engineered microphysiological analysis platform (MAP), designed to model kidney organoids with precision, ensuring uniformity, reproducibility, and rigorous experimental control for pharmaceutical assays. Our innovative kidney organoid MAP employs a biomimetic microfluidic design to supply an organoid array with consistently maintained, balanced extrinsic and intrinsic biochemical conditions. Maintenance of a long-term consistent biochemical balance is a great advantage to precisely regulating organoid development within a recipe that relies on minimal extrinsic factors and intrinsic intercellular communication. In a platform format, the longitudinal and horizontal arrangement of the functional unit, consisting of an organoid chamber, perfusion channels, and interconnection nanochannels, enables the execution of biochemically independent multiplexed experiments within homogeneous arrays of kidney organoids on a single platform. This configuration enhances the systematic design and statistical analysis of organoid experiments.

The MAP’s on-chip organogenesis provides valuable insights into kidney development that might otherwise go unnoticed in other organoid culture formats. The lower perfusion rate required at the CHIR priming on NPC is one representative case (Fig. 2a). Given that higher perfusion rates likely enhance the prevalence of canonical Wnt/β-catenin signaling with increased CHIR99021 flux into the chamber, the probable mechanism behind the developmental success at the lower perfusion rate is the engagement of secondary signaling pathways for NPC development into the nephron. Previous studies suggest the crucial roles of non-canonical Wnt signaling, particularly through Wnt4 autocrine/paracrine signaling^3,37^, which may be compromised under higher perfusion conditions.

The MAP’s nephrogenesis offers promising potential for enhanced controllability and fine-tuning in organoid modeling. For example, the ROCK inhibitor treatment as an effective modulator of kidney organogenesis in MAP presents yet another intriguing on-chip demonstration of signaling dynamics. This outcome suggests that ROCK inhibition, applied through MAP perfusion, serves as a precise signaling to maintain or expand a subset of the NPC and stromal cell pools during Wnt induction^15,16^, which will be subsequently engaged in tubular elongation, especially in the fulfillment of distal segments^38^. Conversely, the proximal segment enrichment prolonged ROCK inhibition underscores the effective guidance of MAP organogenesis, proficiently recruiting the NPC pool to the proximal epithelial construct. This, in turn, underscores another biological aspect of the limited proliferation of proximal tubular epithelial cells^39^ even in the early developing stage, as months-long maturation remained the proximal-enriched pattern.

Our organoid assays highlight the pharmaceutical potential of MAP and the versatility of organoid models for comprehensive analysis of diseases and drug responses. Our broad assessment of multifactorial nephron toxicities provides an efficient and quantitative alternative to traditional qualitative or coarse-grading analyses. Notably, whole-organoid analysis minimizes the risk of biased scrutiny involved in large-size biological entities^40^. Moreover, the incorporation of fluidic dynamics ensures a controlled and continuous introduction of toxicants into the microenvironment, significantly enhancing the study’s accuracy in replicating substance transport and establishing a clear dose-response relationship^41^. Future studies aimed at addressing physiological aspects currently absent, such as ultrafiltration, tubular flows, immune cell interactions, and evaluation of detailed mechanisms of cell death will further enhance organoid responses, allowing for a more accurate representation and understanding of human physiology within our organoid MAP.

The study of ARPKD disease modeling and drug discovery using MAP demonstrates a promising strategy for addressing this incurable infantile polycystic kidney disease (PKD) or other cystic disease, including the more common autosomal dominant PKD. Our organoid modeling, enhanced with CRISPR genetic engineering, provides an effective approach for studying pathological progression, designing therapeutic interventions, and conducting in-depth analyses of biological mechanisms. Notably, the PKHD1-mutated kidney organoids demonstrated spontaneous cystogenesis without requiring external stimulation, such as the intentional elevation of cAMP levels. This characteristic allows for more direct and physiologically relevant molecular comparisons among isogenic models without the induction of pathological phenotypes. In our pursuit of safe treatments for fetal or infant patients, our therapeutic approach was developed within a generalized framework focusing on oxidative stress and NF-κB. The therapeutic efficacy, additionally confirmed through a cAMP-stimulated cystogenic condition, was determined by our comparative RNA-seq analysis within the context of the extracellular matrix and cytoskeleton, across multiple screening levels of significant genes. Among the PKD-associated genes^42^, our ARPKD model demonstrated significant alterations in 47 genes, of which 20 genes showed potential improvement with the proposed therapeutics. Collectively, our insights gained through MAP’s disease modeling and drug study, based on advanced standardized and high-throughput organogenesis, have generated a valuable bioinformatic dataset that can be fine-tuned for next-generation drug discovery. In ongoing and future studies, we aspire to extend our findings to create a therapeutic window for early neonates afflicted with the disease by effectively and safely mitigating kidney pathogenesis.

The precision microfluidic engineering offered by MAP in shaping the microenvironment holds significant potential for various organoid models, particularly given the prevalent reliance on minimal-factor designs. We anticipate that MAP’s capabilities will facilitate overcoming current technical challenges and uncertainties in organoid experiments, resulting in a standardized model that remains consistent over time and across different settings. This will further empower the broad promises associated with organoid research. In this regard, our organoid MAP and the dynamic formation of organoids within the physiological microenvironment will evolve into an advanced, scalable, integrative, and automatable platform technology that can parallel the progress of organoid research.

## METHODS

### Preparation of kidney organoid MAP

MAP was assembled from three distinct components, each individually fabricated and then joined together. To produce the microfluidic unit, we optimized our grayscale photolithography to generate a mold with the three-dimensional fluidic geometries, which were then replicated as polymer chips through the polydimethylsiloxane (PDMS) process. While the procedure adhered closely to the methodology outlined in previous literature^43^, minor adjustments were made in the size of the organoid chamber for optimal growth of kidney organoids. The media reservoirs were fabricated by CO_2_ laser machining of 0.5” thick acrylic plates. After laser machining, any residual debris was thoroughly cleaned in water sonication over several days.

A glass slide (12.5cm x 6.5cm) was perforated with holes using diamond-coated bits (diameter=0.7mm) to enable fluidic flows between the media reservoirs and the PDMS chip. The final assembly phase entailed connecting all components through O_2_ plasma bonding, which ensured the firm bonding of the glass and the PDMS chip. Additionally, uncured PDMS bonding was employed to link the glass with the reservoirs. The assembled device was stored in an airtight, sterile container and subjected to UVO treatment (Model No. 42 Jelight) prior to conducting an organoid experiment.

### On-chip kidney organogenesis in MAP

To generate a large-scale pool of nephron progenitor cells (NPC), a human embryonic pluripotent stem cell (WA09 WiCell) was cultured on a Matrigel-coated six-well plate and differentiated into NPC following the previously published protocol^14,44^. By day 9 of differentiation, a significant portion of the population exhibited markers characteristic of metanephric mesenchyme, with over 70% of the cells being SIX2-positive. Subsequently, gentle enzymatic dissociation using Accutase (ThermoFisher Scientific) allowed us to generate 20-30 cryopreservation vials per NPC batch, each containing approximately 2 million NPCs. These vials were then stored in a liquid nitrogen tank until they were needed for MAP experiments.

For loading NPC into the MAP system, cryopreserved cells were rapidly thawed in a warm bath, diluted with basal media containing 10µM Y-27632 (Selleck Chemicals LLC), centrifuged at 100g for 3 minutes, and resuspended in basal media with 10ng·mL^-1^ FGF9 (R&D Systems, Inc) and 10µM Y-27632. The NPC cells, diluted to the desired concentration, were then pipetted into individual arrays of hemispherical chambers.

After eight hours, the NPC aggregates were transferred to a dynamic culture setting using our custom automatic tilting plate. Our programmed angular control system maintained a constant flow rate of 80 µL·day^-1^ per perfusion unit, with the exception of the 2-day CHIR99021 priming phase described in the Results section. The perfusion medium composition following NPC aggregation was as follows: a 2-day perfusion with 3µM CHIR99021 and 10ng·mL^-1^ FGF9, followed by a 3-day perfusion with 10ng·mL^-1^ FGF9. Subsequently, the organoids were maintained under basal media perfusion. The basal media consisted of advanced RPMI (ThermoFisher Scientific), supplemented with 1× Glutamax (ThermoFisher Scientific), and was free of antibiotics.

### Immunostaining and transcriptomic analysis

To investigate the structural details of kidney organoids, the organoids underwent immunostaining during the perfusion in the MAP system. The following serial steps were designed for on-chip fixation with an 8-hour perfusion of 4% paraformaldehyde (PFA): (1) 8-hour perfusion of PBS for PFA washing. (2) 8-hour perfusion of 0.1% triton-X for cell permeabilization. (3) 1-day perfusion of 2% BSA in PBST for blocking. (4) 1-day perfusion of primary antibodies in 1% BSA in PBST. (5) 1-day perfusion in 1% BSA in PBST to wash away unconjugated antibodies. (6) 1-day perfusion of secondary antibodies in 1% BSA in PBST. (7) 1-day perfusions of PBST and PBS. (8) 8-hour perfusion of an optical clearing solution (RapiClear 1.52; SunJin Lab).

The antibodies used in this study included anti-podocalyxin (AF1658, goat, R&D Systems, Inc), anti-E-Cadherin (ab11512, rat, Abcam), biotinylated Lotus tetragonolobus lectin (B-1325-2, Vector Laboratories), and anti-KIM-1 (MAB1750, mouse, R&D Systems, Inc), diluted at concentrations of 10µg·mL^-1^, 5µg·mL^-1^, 5µg·mL^-1^, and 10µg·mL^-1^, respectively. Secondary antibodies were used at a 1:800 dilution to match the target species, and the streptavidin-biotin reaction (MilliporeSigma) was employed. The antibody-stained and optically cleared organoids were imaged on-chip through the top glass substrate using an upright confocal microscope (Zeiss LMS 710).

For apoptotic and dead cell staining, live organoids were perfused with the Caspase-3/7 Green Cytometry Assay Kit (ThermoFisher Scientific) and propidium iodide (ThermoFisher Scientific) for 1 hour. As these assays do not involve fixation, organoid confocal imaging was performed on a 37°C hot plate. To correlate the locations of apoptotic and dead cells, the organoids were subsequently fixed and stained with LTL and DAPI without membrane permeabilization. Given their stable positioning within individual chambers, the first and second imaging sets could be effectively compared.

To extract RNA from the organoids, we perfused Buffer RLT Plus from the RNeasy Plus Mini Kit (Qiagen) through the ports connecting the hemispherical chamber arrays. The individual perfusates, each containing a lysate from ten organoids, underwent further processing using the additional components provided in the RNeasy Plus Mini Kit to purify total RNA, following the manufacturer’s protocol. Individual RNA samples for RNA-seq analysis, which was pooled from 30 organoids developed across three perfusion sets, were prepared from two independent experiments. The purified RNA samples were promptly stored at -80°C and subsequently sent to a commercial RNA sequencing service (Genewiz, Azenta Life Sciences).

The paired-end RNA sequencing data, generated by the service, underwent initial processing steps, which included adapter trimming and quality assessment using Trim Galore (v0.6.10, Babraham Bioinformatics). The resulting trimmed fastq files were subsequently imported into the RNA-Seq Alignment App within BaseSpace (Illumina). The alignment was performed using the STAR aligner and the human genome reference GRCh38. Significant differences in transcript abundance were assessed using the DESeq2 application within BaseSpace, with a q-value threshold set at 0.05.

A series of gene ontology analyses were performed using the web-based platform, ExpressAnalyst^45^, to gain insights into the functions and attributes of differentially expressed genes. For each analysis, official gene symbols and log2(fold change) values were provided, and the analyses were conducted across four main domains: Biological Process, Molecular Function, Cellular Component, and Reactome pathways. The selection of the most statistically significant results was based on p-values.

### Nephrotoxicity study of kidney organoid MAP

The two drugs under study, cisplatin and gentamicin, were sourced from Millipore Sigma and ThermoFisher Scientific, respectively. For the dose-dependent study, a 21.6mM gentamicin solution was serially diluted in basal media to obtain concentrations of 100, 50, 20, and 10 µM. A cisplatin stock solution was prepared at 20mM in DMSO, and studied concentrations were created through serial dilution in basal media to reach 20, 10, 5, and 2 µM. The control with 0 µM cisplatin served as a vehicle control with the DMSO content in the 20 µM case. Media containing the drug test concentrations were prewarmed and loaded into the media reservoirs to initiate perfusion. For post-cisplatin characterization, assessing KIM-1 and apoptotic/dead cell expression, the media reservoirs were rinsed three times with basal media and then refilled with prewarmed fresh basal media.

To assess nephrotoxicity in the organoids, z-sectioned confocal images were imported into ImageJ and overlaid with a 30×30 µm grid. Guided by this grid and utilizing 18 µm thick z-projections, the biological features within individual microscale partitions were categorized into nephron segment-specific contents, including podocalyxin-positive glomerulus, LTL-positive proximal tubule, and E-Cadherin-positive distal tubule. Additionally, parameters such as KIM-1 positivity, tubular obstruction, abnormal nuclear morphology, presence of caspase 3/7 activity, and PI-positive dead cells were evaluated. Each count was subsequently normalized based on the total number of partitions within the organoid, enabling the assessment of the condition of each organoid.

### Therapeutic analysis of ARPKD kidney organoids in MAP

To evaluate the therapeutic hypothesis for mitigating the cystogenic development of ARPKD organoids, we acquired parthenolide from Selleck Chemicals LLC, methylene blue from Millipore Sigma, and forskolin also from Millipore Sigma. The stock solutions, prepared according to the manufacturer’s instructions, were serially diluted in the basal media, and administered via MAP perfusion in conjunction with the time-specific required factors. For experiments involving methylene blue, precautions were taken to minimize light exposure. Optical imaging was performed using a yellow-filtered light source to prevent undesired photodegradation.

Given the noticeable cystic development in ARPKD organoids, we quantified the frequency of cystogenesis by counting the number of organoids exhibiting the pathological phenotype. For the study on cAMP-enhanced cystogenesis, a one-day application of parthenolide and methylene blue was followed by a seven-day application of methylene blue, preceding a two-day application of forskolin. This sequence was executed across 25 combinations of methylene blue and forskolin. These combinations were evaluated within seven sets of eight-multiplexed MAP organoid cultures, with a morphological assessment conducted at the conclusion of the two-day forskolin applications.

### Computational simulation

To gain insights into the fluidic dynamics of MAP perfusion, we employed a computational simulation using COMSOL Multiphysics (version 5.2). The simulation focused on a three-dimensional model that represented the functional unit of the organoid MAP. This model included a hemispherical chamber, two perfusion channels, and interconnecting nanochannels.

Two key physical disciplines were integrated into the simulation: fluidic dynamics, which were governed by the input perfusion rate, and molecular transport originating from a hypothetical spheroid’s secretion of a pseudo-molecule. The inlet and outlet boundaries of the perfusion channels were configured to replicate laminar inflow, free of the pseudo-molecule. Our objective was to calculate the relative abundance of the pseudo-molecule secretion under steady-state conditions.

## Statistical Analysis

All statistical tests performed are described in the figure legends. Sample sizes are provided in the figure legends. A two-tailed unpaired Student’s *t*-test assuming unequal variances was conducted to statistically analyze differences between two groups. Microsoft Excel was used to perform statistical analysis. All *p* values are noted in the figure legends and raw data.

## Acknowledgments

This work is supported by grant no. NIH NCAT 5UH3TR002155-0505, R01DK072381 and R37DK39773. We thank P. Kim for technical assistance.

## Author Contributions

L.P.L. proposed and supervised the project. S.H., M.S., and L.P.L. designed the study. S.H. and M.S. performed experiments for data and wrote the manuscript. T.M., R.M., and J.V.B. generated nephron progenitor cells from a human embryonic stem cell. All the authors commented on the manuscript.

