## Supplementary Information for "Dynamic Kidney Organoid Microphysiological Analysis Platform"

**CONTENTS**

**- Supplementary Figure 1. ExpressAnalyst of therapeutic effective gene set.**

**- Supplementary Table 1. Genes associated in the ridgeline plot of PKHD1-caused expression difference.**

**- Supplementary Table 2. Genes associated in ridgeline plot of the therapeutics-causing expression difference.**

**Supplementary Figure 1. ExpressAnalyst of therapeutic effective gene set.** The gene ontology and associated genes shown in Fig. 4i are detailed in this figure.

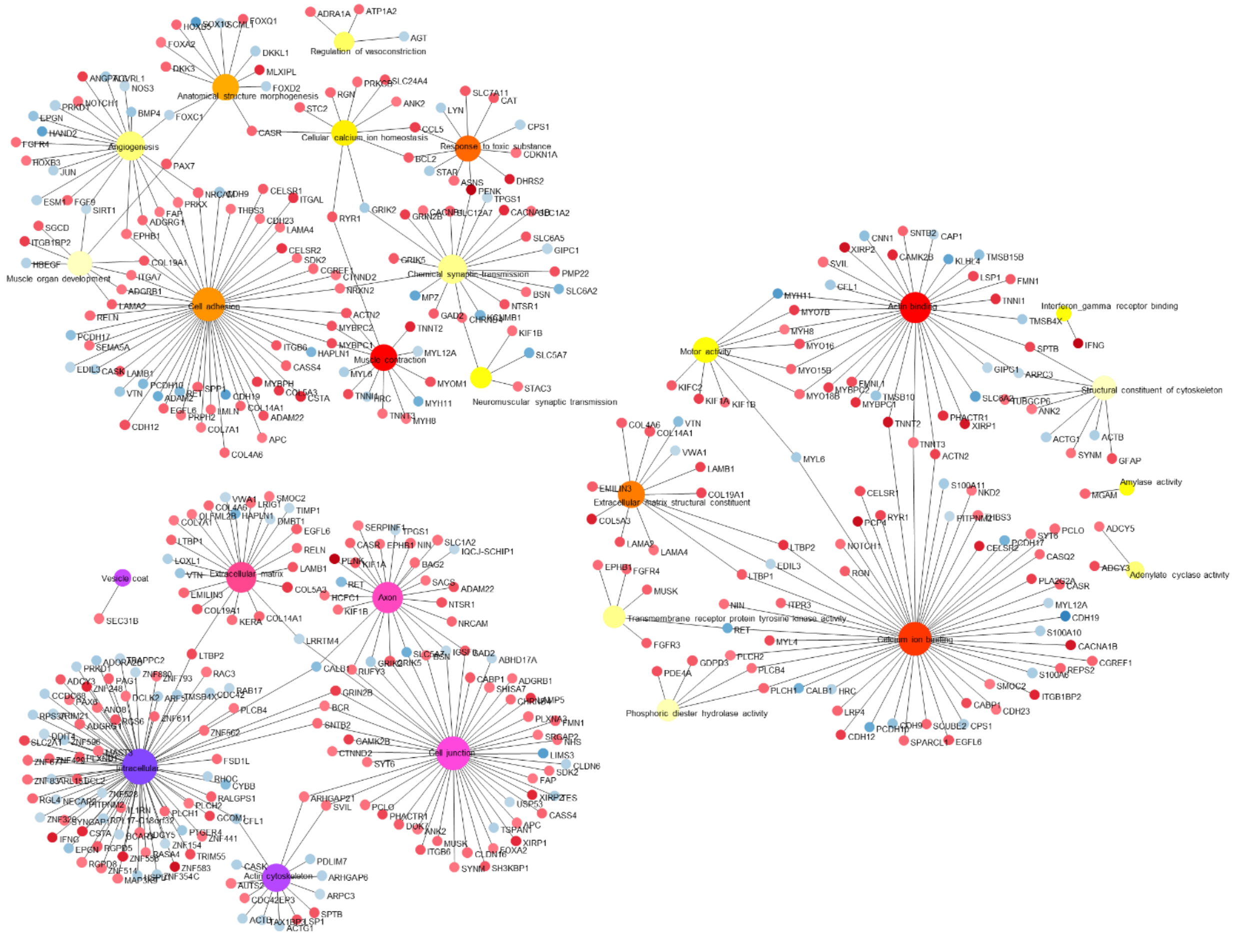

**Supplementary Table 1. Genes associated in the ridgeline plot of PKHD1-caused expression difference.** The list shows gene information shown in Fig. 4g.

| Biological process Skeletal muscle dev. | | Molecular Function Struc. const. of muscle | | Cellular component Protein. ext. matrix | | Reactome Striated Muscle Cont. | |
|---|---|---|---|---|---|---|---|
| Gene | log2(fold) | Gene | log2(fold) | Gene | log2(fold) | Gene | log2(fold) |
| VAMP5 | 2.7 | MYH11 | 1.6 | SBSPON | 2.12 | TPM2 | -0.64 |
| AVPR1A | 1.64 | MYL9 | 0.79 | DMBT1 | 2.11 | DES | -0.81 |
| TSC22D3 | 1.15 | MYL6 | 0.5 | CHI3L1 | 2.07 | NEB | -1.16 |
| NPHS1 | 1.1 | NEBL | 0.44 | GLDN | 1.93 | TNNT2 | -1.52 |
| TCF21 | 0.94 | TPM2 | -0.64 | SOST | 1.74 | TNNI1 | -2.3 |
| CFLAR | 0.77 | SYNM | -0.95 | WNT9B | 1.52 | TNNT3 | -2.92 |
| IGFBP3 | 0.75 | NEXN | -1.07 | TFPI2 | 1.3 | MYBPC2 | -3.17 |
| TGFB1 | 0.74 | NEB | -1.16 | C1QTNF3 | 1.27 | MYL4 | -3.71 |
| F2R | 0.72 | MYOM1 | -1.67 | HAPLN1 | 1.22 | TNNT1 | -3.87 |
| GPHN | 0.61 | JPH1 | -1.85 | FGF1 | 1.21 | TNNC1 | -4.02 |

| | | | | | | | |
|---|---|---|---|---|---|---|---|
| BMP4 | 0.54 | MYBPC2 | -3.17 | SNCA | 1.1 | TNNI2 | -5.37 |
| MYL6 | 0.5 | MYL4 | -3.71 | SPOCK2 | 0.93 | ACTN2 | -5.61 |
| BCL2 | -0.47 | PDLIM3 | -3.79 | ADAMTS19 | 0.93 | MYH3 | -5.71 |
| PDZRN3 | -0.47 | TTN | -3.83 | ADAMTS6 | 0.9 | ACTC1 | -5.74 |
| SKI | -0.54 | MYOT | -4.8 | TGFBR3 | 0.79 | TNNC2 | -5.81 |
| DDX17 | -0.57 | ACTN2 | -5.61 | TINAG | 0.77 | ACTA1 | -5.84 |
| ZBTB18 | -0.6 | MYL1 | -6.18 | NTN4 | 0.75 | MYL1 | -6.18 |
| CACNA2D2 | -0.63 | MYLPF | -6.32 | TGFB1 | 0.74 | MYH8 | -6.85 |
| DMPK | -0.64 | MYH8 | -6.85 | CRTAC1 | 0.69 | MYBPC1 | -6.88 |
| HOXD10 | -0.67 | MYBPC1 | -6.88 | FLRT3 | 0.67 | | |
| ZFHX3 | -0.78 | MYBPH | -7.6 | EFEMP1 | 0.66 | | |
| HDAC9 | -0.81 | | | ANOS1 | 0.66 | | |
| MAMSTR | -0.9 | | | EMID1 | 0.63 | | |
| HLX | -0.92 | | | TIMP1 | 0.61 | | |
| SIX4 | -0.92 | | | CASK | 0.58 | | |
| SOX8 | -0.93 | | | BMP4 | 0.54 | | |
| COL19A1 | -0.93 | | | VWA1 | 0.48 | | |
| SIX1 | -0.94 | | | CST3 | 0.48 | | |
| ADAM12 | -0.94 | | | NPNT | 0.47 | | |
| ITGA7 | -0.98 | | | COL26A1 | -0.45 | | |
| BOC | -1.02 | | | NAV2 | -0.45 | | |
| NOTCH1 | -1.05 | | | LAMA1 | -0.51 | | |
| DOK7 | -1.08 | | | LAMB1 | -0.56 | | |
| MEOX2 | -1.12 | | | COL4A6 | -0.56 | | |
| MEF2C | -1.16 | | | LTBP4 | -0.56 | | |
| RXRG | -1.18 | | | ADAMTS10 | -0.58 | | |
| ELN | -1.19 | | | ADAMTS2 | -0.66 | | |
| LRP4 | -1.2 | | | SPON1 | -0.69 | | |
| SVIL | -1.2 | | | VWC2 | -0.69 | | |
| MEGF10 | -1.21 | | | LAMA2 | -0.7 | | |
| NUPR1 | -1.36 | | | COL23A1 | -0.71 | | |
| CDON | -1.39 | | | LTBP2 | -0.71 | | |
| DNER | -1.43 | | | ADAMTS12 | -0.72 | | |
| MSTN | -1.49 | | | COL5A1 | -0.75 | | |
| WNT3A | -1.79 | | | COL27A1 | -0.77 | | |
| PITX2 | -1.83 | | | SPOCK3 | -0.8 | | |
| WNT10B | -2.07 | | | C1QTNF5 | -0.85 | | |
| MYF6 | -2.17 | | | LAMA4 | -0.88 | | |
| UNC13A | -2.27 | | | TNXB | -0.88 | | |
| RBM24 | -2.34 | | | NID2 | -0.93 | | |
| IGF1 | -2.37 | | | COL19A1 | -0.93 | | |
| PITX1 | -2.71 | | | ABI3BP | -0.96 | | |
| RYR1 | -3.17 | | | COL21A1 | -1.02 | | |
| SRPK3 | -3.44 | | | FBN3 | -1.03 | | |
| NMRK2 | -3.81 | | | MGP | -1.03 | | |
| CACNG2 | -3.88 | | | COL8A2 | -1.04 | | |
| MUSK | -4.2 | | | SMOC2 | -1.05 | | |
| CHRNA1 | -4.24 | | | RUNX1 | -1.1 | | |
| CACNA1S | -4.41 | | | ACHE | -1.11 | | |
| CAV3 | -4.72 | | | LTBP1 | -1.14 | | |
| CHRND | -5.36 | | | WNT11 | -1.16 | | |
| KLHL41 | -5.44 | | | GPC4 | -1.18 | | |
| VGLL2 | -5.44 | | | CHL1 | -1.18 | | |
| MYOG | -5.61 | | | ELN | -1.19 | | |
| MYOD1 | -5.72 | | | FGF9 | -1.19 | | |
| SMYD1 | -5.79 | | | EGFL6 | -1.19 | | |

| | | | | | |
|---|---|---|---|---|---|
| ACTA1 | -5.84 | | | C1QL1 | -1.2 |
| PAX7 | -6.13 | | | EGFLAM | -1.22 |
| MYLPF | -6.32 | | | COL7A1 | -1.22 |
| MYF5 | -6.82 | | | COL6A6 | -1.24 |
| | | | | LGALS3 | -1.26 |
| | | | | COL22A1 | -1.29 |
| | | | | CILP2 | -1.39 |
| | | | | COL14A1 | -1.41 |
| | | | | ADAMTS14 | -1.42 |
| | | | | WNT3 | -1.58 |
| | | | | BCAN | -1.59 |
| | | | | LOX | -1.6 |
| | | | | EMILIN3 | -1.6 |
| | | | | ADAMTS18 | -1.6 |
| | | | | ADAMTS17 | -1.63 |
| | | | | GPC2 | -1.64 |
| | | | | RELN | -1.66 |
| | | | | COL12A1 | -1.67 |
| | | | | THSD4 | -1.68 |
| | | | | PODN | -1.71 |
| | | | | CTHRC1 | -1.72 |
| | | | | WNT9A | -1.72 |
| | | | | WNT2B | -1.77 |
| | | | | SPOCK1 | -1.78 |
| | | | | WNT3A | -1.79 |
| | | | | COCH | -1.88 |
| | | | | THBS2 | -1.98 |
| | | | | NTN1 | -2 |
| | | | | OGN | -2.02 |
| | | | | WNT10B | -2.07 |
| | | | | LAMA3 | -2.13 |
| | | | | SPARCL1 | -2.16 |
| | | | | SLC1A3 | -2.4 |
| | | | | PTPRZ1 | -2.51 |
| | | | | ITGB4 | -2.65 |
| | | | | COL5A3 | -2.72 |
| | | | | LUM | -2.98 |
| | | | | KERA | -3.23 |
| | | | | WNT7A | -4.08 |
| | | | | ASPN | -4.8 |

**Supplementary Table 2. Genes associated in ridgeline plot of the therapeutics-causing expression difference.** The list shows gene information shown in Fig. 4h

| **Biological process** | | **Molecular Function** | | **Cellular component** | | **Reactome** | |
|---|---|---|---|---|---|---|---|
| **Ext. structure org.** | | **Collagen binding** | | **Protein. ext. matrix** | | **HS-GAG degradation** | |
| Gene | log2(fold) | Gene | log2(fold) | Gene | log2(fold) | Gene | log2(fold) |
| COL9A1 | -1.86 | MMP9 | -1.68 | MATN1 | -2.66 | GPC5 | -1.09 |
| VMO1 | -1.71 | ITGA10 | -1.55 | MMP17 | -1.92 | SDC1 | -0.9 |
| MMP9 | -1.68 | PODN | -1.47 | COL9A1 | -1.86 | GPC2 | -0.89 |
| RGCC | -1.55 | CD44 | -0.89 | MMP9 | -1.68 | GPC1 | -0.5 |
| SMOC1 | -1.53 | PCOLCE | -0.69 | HAPLN3 | -1.53 | SDC3 | -0.31 |
| COL8A1 | -1.5 | ITGA11 | -0.64 | SMOC1 | -1.53 | SDC2 | 0.37 |

| | |
|---|---|
| COL8A2 | -1.43 |
| EGFLAM | -1.33 |
| GREM1 | -1.31 |
| LGALS3 | -1.3 |
| GPM6B | -1.27 |
| CSGALNACT1 | -1.22 |
| COL21A1 | -1.17 |
| LOXL2 | -1.16 |
| ENG | -1.12 |
| COL9A3 | -1.09 |
| ADAMTS2 | -0.99 |
| RAMP2 | -0.91 |
| COL15A1 | -0.88 |
| SOX9 | -0.86 |
| COL24A1 | -0.85 |
| THSD4 | -0.83 |
| KAZALD1 | -0.78 |
| TFAP2A | -0.77 |
| TIMP2 | -0.74 |
| MMP2 | -0.72 |
| TGFB2 | -0.68 |
| COL26A1 | -0.66 |
| COL6A2 | -0.64 |
| COL5A1 | -0.64 |
| SULF2 | -0.62 |
| TGFBI | -0.61 |
| SFRP2 | -0.6 |
| FBLN1 | -0.53 |
| SH3PXD2B | -0.53 |
| POSTN | -0.49 |
| DDR2 | -0.48 |
| PDGFRA | -0.48 |
| CCDC80 | -0.41 |
| SERPINH1 | -0.39 |
| COL4A5 | 0.35 |
| CST3 | 0.38 |
| PXDN | 0.38 |
| SERPINF2 | 0.4 |
| COL4A1 | 0.4 |
| COL4A6 | 0.41 |
| P3H3 | 0.43 |
| APLP2 | 0.45 |
| COL14A1 | 0.5 |
| HPN | 0.51 |
| EGFL6 | 0.53 |
| MMP14 | 0.57 |
| LAMB2 | 0.61 |
| DPP4 | 0.63 |
| NID1 | 0.63 |
| COL4A2 | 0.73 |
| COL18A1 | 0.74 |
| FOXC2 | 0.78 |
| LAMC1 | 0.79 |
| COL4A4 | 0.8 |
| NPNT | 0.81 |
| COLGALT2 | 0.9 |

| | |
|---|---|
| DDR2 | -0.48 |
| FN1 | -0.44 |
| SERPINH1 | -0.39 |
| PAK1 | 0.31 |
| ADAM9 | 0.5 |
| COL14A1 | 0.5 |
| NID1 | 0.63 |
| SPARC | 0.66 |
| ITGA1 | 0.81 |
| PDGFA | 0.88 |
| DDR1 | 0.96 |
| ECM2 | 1.4 |
| NID2 | 1.81 |

| | |
|---|---|
| COL8A1 | -1.5 |
| PTPRZ1 | -1.48 |
| PODN | -1.47 |
| HAPLN1 | -1.44 |
| COL8A2 | -1.43 |
| VWC2 | -1.37 |
| ADAMTS19 | -1.34 |
| EGFLAM | -1.33 |
| LGALS3 | -1.3 |
| COL21A1 | -1.17 |
| LOXL2 | -1.16 |
| SPOCK3 | -1.11 |
| COL9A3 | -1.09 |
| GPC5 | -1.09 |
| WNT11 | -1.03 |
| NTN1 | -1 |
| ADAMTS2 | -0.99 |
| ADAMTS6 | -0.92 |
| SLC1A3 | -0.91 |
| GPC2 | -0.89 |
| COL15A1 | -0.88 |
| FBN3 | -0.85 |
| COL24A1 | -0.85 |
| THSD4 | -0.83 |
| BMP4 | -0.81 |
| KAZALD1 | -0.78 |
| ADAMTS4 | -0.76 |
| TIMP2 | -0.74 |
| C1QTNF5 | -0.73 |
| SPON2 | -0.73 |
| MMP2 | -0.72 |
| SFRP1 | -0.68 |
| MFAP4 | -0.68 |
| COL26A1 | -0.66 |
| SPON1 | -0.65 |
| COL6A2 | -0.64 |
| COL5A1 | -0.64 |
| TGFBI | -0.61 |
| ADAMTS10 | -0.59 |
| ADAMTS12 | -0.54 |
| FBLN1 | -0.53 |
| SPOCK1 | -0.52 |
| ADAMTS7 | -0.51 |
| GPC1 | -0.5 |
| POSTN | -0.49 |
| CD248 | -0.47 |
| FN1 | -0.44 |
| CCDC80 | -0.41 |
| MFAP2 | -0.39 |
| CASK | -0.32 |
| FLRT3 | 0.27 |
| COL4A5 | 0.35 |
| CST3 | 0.38 |
| PXDN | 0.38 |
| COL4A1 | 0.4 |
| COL4A6 | 0.41 |

| | |
|---|---|
| GUSB | 0.42 |
| SGSH | 0.48 |
| GPC6 | 0.52 |
| GPC4 | 0.54 |
| GLB1 | 0.67 |
| AGRN | 0.77 |
| NAGLU | 1.05 |

| | |
|---|---|
| ADAMTSL4 | 0.91 |
| DDR1 | 0.96 |
| MMP1 | 0.97 |
| P3H2 | 0.99 |
| PLOD2 | 1.03 |
| SNCA | 1.18 |
| SPOCK2 | 1.22 |
| WT1 | 1.25 |
| GAS6 | 1.27 |
| ECM2 | 1.4 |
| NPHS1 | 1.66 |
| OLFML2A | 1.76 |

| | |
|---|---|
| ERBIN | 0.42 |
| EFEMP1 | 0.49 |
| LAMC3 | 0.5 |
| COL14A1 | 0.5 |
| VEGFA | 0.51 |
| GPC6 | 0.52 |
| EGFL6 | 0.53 |
| GPC4 | 0.54 |
| ADAMTS9 | 0.55 |
| DAG1 | 0.58 |
| LAMA2 | 0.59 |
| LAMB2 | 0.61 |
| NTN4 | 0.63 |
| NID1 | 0.63 |
| C1QL4 | 0.63 |
| LGALS3BP | 0.66 |
| SPARC | 0.66 |
| TFPI2 | 0.66 |
| FREM1 | 0.68 |
| COL4A2 | 0.73 |
| COL18A1 | 0.74 |
| ENTPD2 | 0.74 |
| TGFBR3 | 0.76 |
| AGRN | 0.77 |
| LAMB1 | 0.77 |
| LAMC1 | 0.79 |
| COLEC12 | 0.79 |
| COL4A4 | 0.8 |
| NPNT | 0.81 |
| ADAMTS1 | 0.85 |
| ADAMTS15 | 0.86 |
| MAMDC2 | 0.86 |
| SOST | 0.91 |
| ADAMTSL4 | 0.91 |
| CILP | 0.95 |
| CCBE1 | 0.97 |
| MMP1 | 0.97 |
| NAV2 | 0.98 |
| P3H2 | 0.99 |
| LTBP2 | 1.02 |
| LAMA1 | 1.08 |
| FREM2 | 1.09 |
| SNCA | 1.18 |
| SPOCK2 | 1.22 |
| ECM2 | 1.4 |
| FBLN2 | 1.44 |
| CILP2 | 1.52 |
| LAMA5 | 1.53 |
| SPN | 1.69 |
| TFF3 | 1.76 |
| NID2 | 1.81 |
| EMID1 | 1.82 |
| C1QTNF7 | 1.83 |
| EMILIN2 | 2 |
| CHI3L1 | 2.12 |
| CPZ | 2.72 |

| | | | | |
|---|---|---|---|---|
| | | | | CRTAC1 3.55 |
